# Adaptations to Oral and Pharyngeal Swallowing Function Induced by Injury to the Mylohyoid Muscle

**DOI:** 10.1101/832147

**Authors:** Suzanne N. King, Brittany Fletcher, Bradley Kimbel, Nicholas Bonomo, Teresa Pitts

## Abstract

Muscle injury is a frequent side effect of radiation treatment for head and neck cancer. To understand the pathophysiology of injury-related dysfunction, we investigated the effects of a single muscle injury to the mylohyoid on oropharyngeal swallowing function in the rat. The mylohyoid protects the airway from food/liquid via hyolaryngeal elevation and plays an active role during both oral and pharyngeal swallowing. We hypothesized (1) that fibrosis to the mylohyoid alters swallowing bolus flow and licking patterns; (2) that injury to the mylohyoid changes normal activity of submental, laryngeal, and pharyngeal muscles during swallowing. A chilled cryoprobe was applied to the rat mylohyoid muscle to create a localized injury. After 1- and 2-weeks post-injury, swallowing bolus transit was assessed via videofluoroscopy and licking behavior via an electrical lick sensor. The motor activity of five swallow-related muscles were analyzed immediately after injury using electromyography (EMG). Comparisons were made pre- and post-injury. Fibrosis was confirmed in the mylohyoid at 2-weeks post-injury by measuring collagen content. One-week after injury, bolus size decreased, swallowing rate reduced, and licking patterns were altered. Immediately post-injury, there was a significant depression in mylohyoid and thyropharyngeus EMG amplitudes during swallowing. Our results demonstrated that injury to the mylohyoid is sufficient to cause changes in deglutition. These disruptions in oral and pharyngeal swallowing were detected prior to long-term fibrotic changes, including delays in tongue movement, alterations in bolus flow, and changes in sensorimotor function. Therefore, injuring a single important swallowing muscle can have dramatic clinical effects.

## Introduction

Dysphagia is a major complication of radiation treatment for head and neck cancer. Radiation associated dysphagia includes reductions in base of tongue retraction, hyolaryngeal displacement, and movement of pharyngeal structures [1-4]. This leads to prolonged oral and pharyngeal transit times, as well as incomplete clearance of saliva and food materials from within the pharynx [3,4]. As a result, patients often require long-term diet modifications, placement of feeding tube to maintain nutrition, and/or behavior modifications to prevent aspiration [5,6]. Clinical reports suggest that muscle injury and/or fibrosis is a driving mechanism behind these swallowing deficits, restricting mobility [7-9]. One approach to limiting these devastating impairments is to spare dysphagia-related structures, and thus, reduce the severity of the injury. However, identifying at-risk organs is complicated in humans because swallowing consists of several integrated movements generated by multiple muscles [10-13]. While previous animal research has focused on studying the effects of radiation injuries [14,15], it is difficult to elucidate the individual impact one muscle has on contributing to dysphagia from previous work. This is important because injury to a single muscle may contribute to movement dysfunction via structural and/or neuroplastic changes. Further research is needed to understand muscle specific injury adaptations associated with swallowing dysfunction. This information is critical to understanding the pathophysiology of dysphagia associated with muscle injury.

Recent evidence has shown that the submental muscles (e.g., the mylohyoid) are at risk for developing radiation-associated dysphagia [13,16]. Mylohyoid activity is tightly regulated, contributing to both the oral and pharyngeal phases of swallowing [17-19]. During the oral phase, it aids in mandibular depression and tongue elevation, which facilitates chewing and the propulsion of food posteriorly through the oropharynx during swallowing [17,20]. During pharyngeal swallowing, it plays a vital role as the leading complex of muscles recruited at the onset of swallowing [18]. Contraction of the mylohyoid raises the hyolaryngeal complex, which causes deflection of the epiglottis to protect the airway and aids in opening the upper esophageal sphincter to allow food to enter into the esophagus [21]. The multiple functions performed by the mylohyoid suggest that an injury to this muscle may have a devastating impact on deglutition.

In other models of muscle injury, disturbances in motor output is driven by changes in neural control either due to a dysfunction of the motor neurons or an alteration in sensory input driving motor output. Sensory input is critical to motor execution of swallowing and disruptions have been shown to impair the coordination of the oropharyngeal structures [22,23]. The trigeminal nerves (CN V) innervating the mylohyoid muscle have extensive projections to the neural centers controlling the laryngeal and pharyngeal structures involved in swallowing [24,25]. Previous work has demonstrated that capsaicin to the orofacial muscles, i.e. masseter (also innervated by CN V), provokes excitability of neurons in the nucleus tractus solitarius of the dorsal medulla, which is an important area mediating swallowing biomechanics [26,27]. Capsaicin is known to activate nociceptors, which are a subset of sensory neuron that detects tissue damage. This work provides evidence that nociceptive sensory input from one muscle may have widespread effects on the sensorimotor system. This in turn suggests that an injury to the mylohyoid muscle may result in sensory adaptations, which affect the normal function of other swallowing muscles not directly injured.

The objective of this study was to investigate the effects of mylohyoid muscle injury on oropharyngeal swallowing function. We first examined whether mylohyoid injury resulted in deviations in the licking pattern and bolus transit during swallowing using videofluoroscopy and electrophysiological recordings of self-drinking rats. We hypothesized that injury provokes fibrosis within the mylohyoid, resulting in delays in swallowing bolus flow and alterations in rhythmic licking behaviors. Next, utilizing an anesthetized animal model, we determined if swallowing dysfunction after mylohyoid injury is purely a mechanical problem or if it is a sensory mediated event. The submental (mylohyoid, geniohyoid), laryngeal (thyroarytenoid, thyrohyoid), and pharyngeal (thyropharyngeus) muscles, which are vital to pharyngeal swallowing are innervated by different cranial and spinal nerves. We hypothesized that swallowing dysfunction after mylohyoid injury results in changes in the normal motor activity of swallowing muscles not directly injured.

## Methods

Eleven male Sprague Dawley rats (450-500g; ∼8.5 - 9 months old) were used for these studies. All experimental protocols were approved by the Institutional Animal Care and Use Committee of the University of Louisville.

### Mylohyoid Cryoinjury Procedure

To injure the mylohyoid, we employed a cryoinjury model to inflict tissue damage similar to that caused by radiation injury. Cryoinjury is a commonly used model for inducing muscle fibrosis in the heart [28,29], bladder [30], hindlimb [31], and others [32]. The method applies a liquid nitrogen-cooled cryoprobe to the muscle of interest. This generates local tissue damage in relatively uniform size and shape to control for severity and specificity of the injury to only the mylohyoid muscle. Radiation-induced injuries are thought to be provoked by microvascular damage and persistent oxidative stress, gradually leading to excessive accumulation of collagen and aberrant extracellular matrix formation [33]. The molecular mechanisms of cryoinjury are similar to those caused by irradiation, specifically with the occurrence of prominent inflammation and necrosis, ending with the formation of fibrosis within 2 weeks [34].

Six rats underwent bilateral cryoinjuries to the belly of the mylohyoid muscles and were followed for 1- and 2-weeks post-injury to assess lick and swallowing function. Animals were initially anesthetized with isoflurane (2-3%) via inhalation and then transitioned to Ketamine (90mg/kg) and Xylazine (9mg/kg) with Atropine (0.05mg/kg) via intraperitoneal (IP) injection. Buprenorphine 0.01–0.05mg/kg was provided for analgesia for 3-days post. The mylohyoid muscle was surgically exposed via longitudinal incision and blunt dissection at midline of the anterior digastric muscle. The flat end of a cryoprobe (2mm in diameter; Cry-AC-3 B-800, Brymill Cryogenic Systems; UK) was chilled by infusion of liquid nitrogen and applied to the surface of the exposed muscle for 30 seconds. The probe was then non-traumatically detached by allowing it to equilibrate to room temperature prior to removal to prevent further injury to the muscle. Following detachment, a second application in the contralateral muscle belly was applied for another 30 seconds. The incision was then sutured closed. After 2-weeks post injury, animals were euthanized with IP injection of urethane and transcardially perfused with PBS and 4% paraformaldehyde (PFA). Mylohyoid muscles were collected for histological analysis. For normal controls for histology, muscles were harvested from two uninjured rats.

### Histology

To confirm fibrosis in the mylohyoid muscle, tissue was post-fixed in 4% PFA, sucrose cryoprotected, and then embedded in freezing media (OCT compound, Tissue-Tek, CA). Muscle samples were then frozen using an isopentane-liquid nitrogen approach. Serial cross sections were taken with the left muscle, and longitudinal sections were obtained with the right muscle at 8μm via cryostat. Tissue specimens were first stained with Hematoxilin and Eosin to visualize general morphologic changes and denote the lesion site. Presence of inflammatory infiltrate, edema, myonecrosis, and angiogenesis were evaluated. Additional sections were stained with Trichrome Stain, Gomori One-Step, Aniline Blue (Newcomer Supply, Middleton, WI) following manufactures description. Stained slides were examined with a Nikon TiE inverted microscope with Nikon Elements Advanced Research software (Nikon Corp, Melville, NY). Images were analyzed with ImageJ [35] using Fiji plugin. Collagen content was quantified by calculating the relative total area of blue/purple staining within samples. Comparisons were made with two uninjured animals.

### Videofluoroscopic Swallow Study

To quantify changes in bolus flow during swallowing, rats were given chocolate milk (thin consistency) mixed with 40% w/v barium sulfate ad libitum. Animals were acclimated to the testing chamber and testing solution prior to experimental tests. A GE Innova Model 3100 fluoroscope was used at a rate of 30 frames per second. Animals were given 5 minutes to self-feed and 15 second video clips were taken throughout during notable periods of drinking. Videos were identified for analysis if they included more than three uninterrupted, sequential swallows. A minimum of three separate 15 second video clips per animal were randomly chosen and analyzed using ImageJ (NIH, Bethesda, MD). Swallowing events were identified by advancing frame-by-frame. Measurements were taken of 10-15 different swallows for each rat and averaged. A drinking cannula (within field of view) was used as a calibration marker to appropriately scale images for size analysis. The second cervical vertebra was used to denote the end of the pharyngeal swallowing phase. The following VFSS metrics were calculated as previously described [36,37]: jaw excursion rate, inter-swallow interval, lick-swallow ratio, and bolus speed through the pharynx. Bolus area within the pharynx was also calculated for each swallow (mm^2^) by outlining the barium contrast material at the frame where the bolus was positioned in the oropharynx, immediately after retroflexion from the vallecula. Swallow rate is based off the number of swallows that occurred during a period of uninterrupted drinking at the spout.

### Lick Testing

To determine changes in the pattern of rhythmic tongue movements after mylohyoid injury, we recorded licking behaviors using an electrical lick sensor similar to those published previously [38-40]. Rats were trained to self-feed from a custom-designed cage with an opening in the sidewall of the chamber where the drinking spout was located. The shape of the opening is proportional to the rat’s zygomatic arch, to allow for stabilization of the head without restricting jaw movement [38]. A metal spout and metal cage floor were connected to an electrical circuit. Output was generated following tongue contact with the spout, completing the circuit. The stainless steel spout was covered with hard plastic tubing to limit contact of the spout to only the tongue during drinking. Previous work has indicated that lick rate decreases when access to the drinking tube is restrained, which is attributed to the greater lingual effort needed to maintain contact and coordinate movement. Therefore, to obtain measures of true licking ability and limit the ceiling effect, we created a challenging drinking activity by increasing the distance between the rats’ snout and liquid being dispensed from the spout [38]. The distance was kept constant throughout the study to limit environmental influences on licking rhythm. Amplified data was recorded via Spike 2 software (Cambridge Electronic Design, UK). Rats were tested in 10-minute drinking sessions after 3 hours of water deprivation. Training consisted of six trails of drinking within similar testing conditions prior to obtaining baseline data. The following licking variables were calculated pre- and post-injury as previous described [38-42]: contact duration, inter-contact interval, interlick interval, lick frequency, cluster size (duration), number of clusters, and licks per cluster. Interlick intervals (ILI) were calculated as the time between the start of one lick and the onset of a succeeding lick. Only ILIs <500ms were analyzed, which denote anterior/posterior or lateral displacement of the tongue between contact with the spout [39,38,40]. Normal rats ingest liquids in licking clusters, where they perform multiple consecutive licks then take a short break from drinking and then start again. Based on previous reports, a break in licking >500ms in duration indicates that the animal has moved their head away from the spout (this break is called an intercluster interval). The duration of a cluster was calculated as the time between five or more consecutive licks. An intercluster interval of >500ms was used to determine the onset and termination of the licking cluster. This criteria was chosen based off prior research in rats, indicating that the intercluster interval should constitute 2.5x the mode of the ILI [40]. The number of clusters that occurred in a 10-minute period and the number of licks performed within each cluster were also calculated.

### Electromyography (EMG)

Experiments were performed on three spontaneously breathing adult male rats. To determine disruptions in sensorimotor activity during swallowing, we analyzed the EMG signal of five muscles involved in swallowing. Animals were initially anesthetized with isoflurane via inhalation and then transitioned to urethane (1ml/kg; IP). Atropine (0.03mg/kg) was given at the beginning of the experiment to reduce secretions. A cannula was placed in the trachea and body temperature was maintained at 37.5 °C. Bipolar insulated fine wire electrodes were placed into the mylohyoid, geniohyoid, thyrohyoid, thyropharyngeus, and thyroarytenoid muscles as previously described [43,27]. The signal was amplified and recorded at 10 kHz using Power 1401 system (Cambridge Electronic Design, UK) and Spike 2 software. Swallowing was evoked pre- and post-injury using water stimuli by infusing 0.2ml into the pharynx via half-inch polyethylene tubing attached to a 1ml syringe. After obtaining baseline swallows, a cryoinjury was performed unilaterally as described above. Swallows were stimulated approximately every 5 minutes for up to 45 minutes post-injury. At the conclusion of the experiment, animals were euthanized and electrode placement was confirmed. Raw signals were low pass filtered, rectified, and integrated with a time constant of 20ms using Spike 2. Swallows were identified by sequential bursts of mylohyoid and thyrohyoid muscles. Amplitude was measured from EMGs of each muscle and normalized to the percentage of maximum across the experiment. Comparisons were made between pre- and post-injury.

Muscles were selected for this experiment based off their function and diverse innervation. During pharyngeal swallowing the mylohyoid, geniohyoid, and thyrohyoid facilitate anterior-superior hyolaryngeal elevation; thyroarytenoid helps adduct the vocal folds; and thyropharyngeus activity creates a stripping wave pushing the bolus toward the esophagus. These swallowing muscles comprise different cranial and spinal nerves, which permits testing the hypothesis that mylohyoid injury affects the function of muscles not directly injured.

### Statistical analysis

Comparisons were made pre- and post-injury (1- and 2-weeks). Repeated measures analysis of variance was carried out to test main effects for each behavioral assay (licking, swallowing, and EMG). If means differed significantly by *p*-value <0.05, a post hoc Least Squares Means test was performed to compare time effects. T-tests were used to compare differences in collagen content between cryoinjury specimens and uninjured controls. *p*<0.05 was considered statistically significant.

## Results

### Histological Changes to Cryoinjured Mylohyoid Muscle

To confirm that cryoinjury provoked fibrosis within the mylohyoid muscle, we measured the level of collagen content. Significant differences in the percentage of collagen per tissue area were found 2-weeks post-injury (Figure 1). Collagen content was 5.07% higher post-injury compared to uninjured mylohyoid muscles (*p*= 0.004; 95% CI, 1.93 to 8.21). There was minimal inflammatory infiltrate or edema observed in the mylohyoid 2-weeks post-injury.

**Figure 1.**
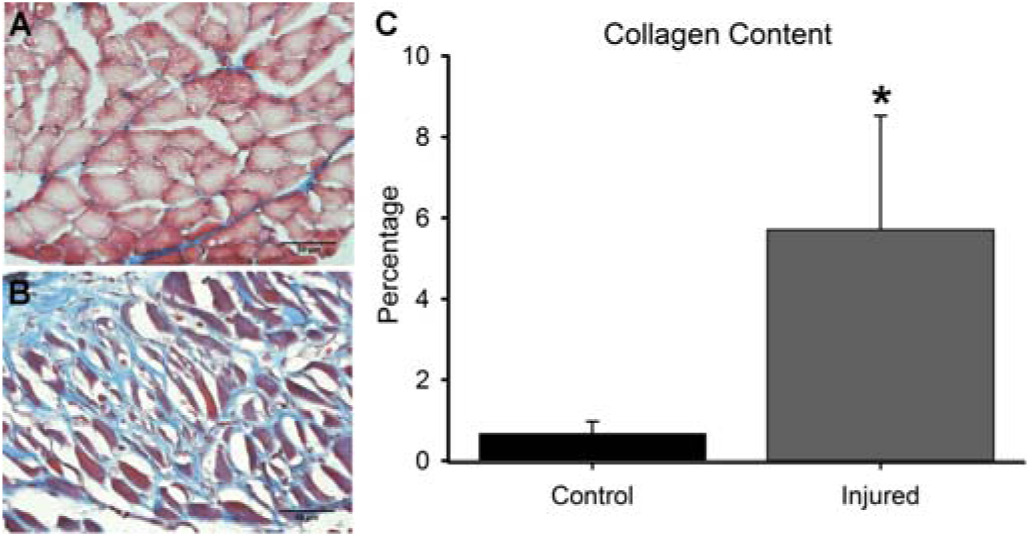
Increased collagen content in the mylohyoid post-injury. Example of a cross section of the mylohyoid muscle from uninjured (A) and injured (B) rats. Blue staining denotes collagen fibers and red/purple indicates muscle (Bar, 50μm). (C) Significant increases in the percentage of collagen was found 2-weeks after cryoinjury compared to non-injured controls (*p* <0.05 is shown as *).

### Videofluoroscopy Swallow Studies

To determine differences in jaw movement and pharyngeal swallowing performance, we compared bolus transit 1- and 2-weeks post-injury to baseline function (Figure 2). Significant differences were found in the jaw excursion rate (cycles/second) after injury (*F*(2,10) = 6.63, *p* = 0.02) with decreases observed 1- and 2-weeks post-injury compared to pre-injury (*p<* 0.03; *p*<0.04). There were significant differences in bolus speed through the pharynx (*F*(2,10) = 7.54, *p* = 0.01). At 1-week post-injury, rats exhibited the slowest bolus speed compared to pre-injury (*p*= 0.02) and 2-weeks post-injury (*p*< 0.04). Bolus area differed significantly after injury (*F*(2,10) = 45.63, *p* = 0.001). Smaller bolus sizes were swallowed 1- and 2-weeks post-injury compared to pre-injury (both *p*< 0.001). Lastly, there were significant differences in swallowing rate (*F*(2,10) = 5.18, *p* = 0.03) with decreases in frequency found 1-week after injury compared to uninjured and 2-weeks post-injury (both *p*< 0.04). No statistically significant differences were found with inter-swallow interval (*F*(2,10) =4.07, *p* = 0.051) and lick-swallow ratio (*F*(2,10) =3.28, *p* = 0.08).

**Figure 2.**
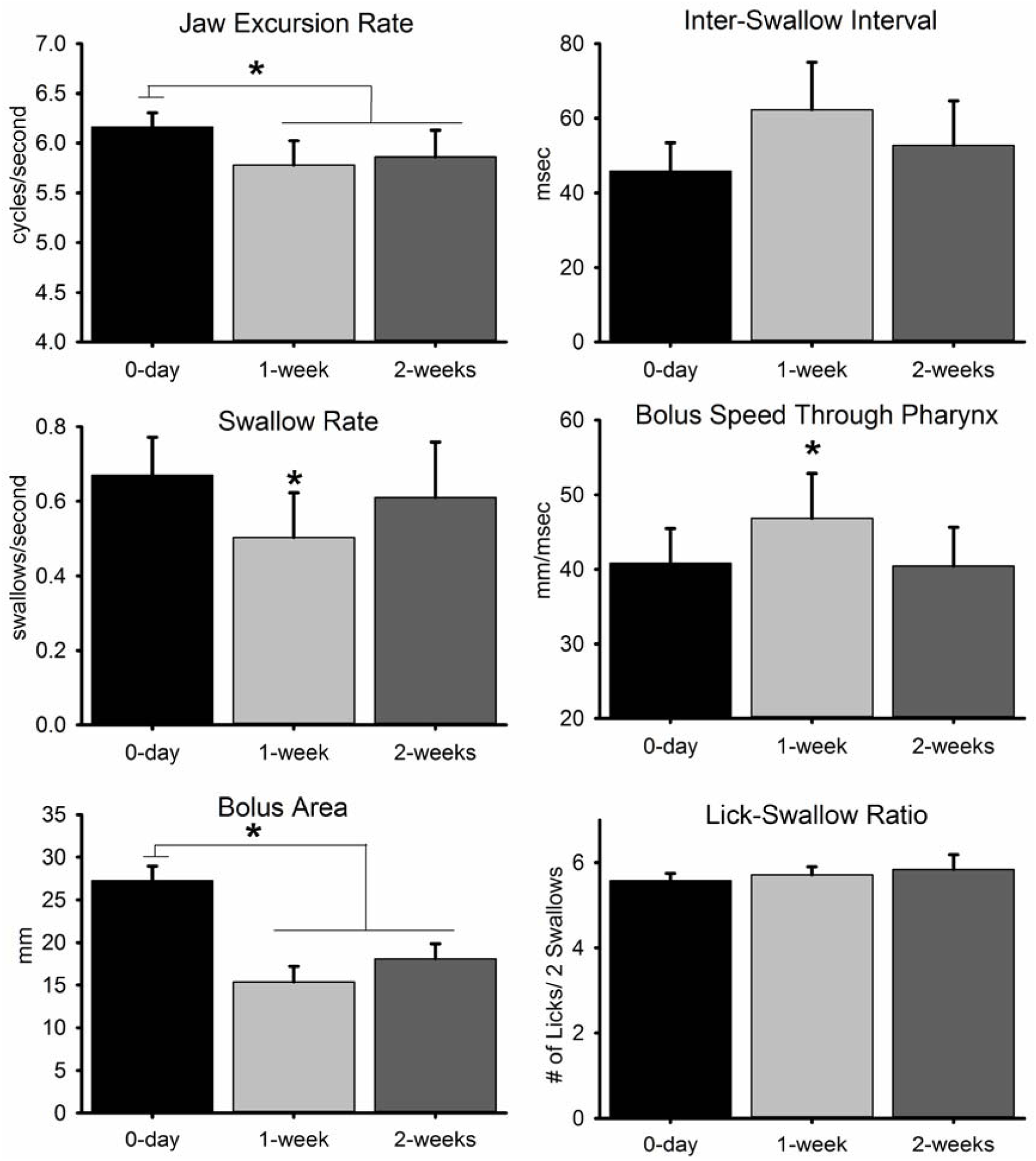
Changes in swallowing metrics taken from videofluoroscopy swallow studies at 0, 1- and 2-weeks post mylohyoid cryoinjury. Results demonstrate the mean and standard deviations for jaw excursion rate, inter-swallow interval, swallow rate, bolus speed through pharynx, bolus area, and lick-swallow ratio. Statistical significance of *p* < 0.05 is indicated by *.

### Alterations in Licking Behaviors

Alterations in the pattern of licking and licking microstructure were found after injury to the mylohyoid muscle (Figure 3 and 4). The total number of licks during the 10-minute drinking session was comparable across time (pre-injury 2075±618 licks; post-injury 1690±463 licks). The average inter-lick interval differed significantly after injury (*F*(2,10) = 10.00, *p* = 0.004) with longer intervals found 1- and 2-weeks post-injury compared to pre-injury (both *p*< 0.01). The average number of licks per second differed across groups (*F*(2,10) = 8.63, *p* = 0.01) with significant decreases after injury compared to pre-injury (1-week: *p*< 0.02; 2-weeks: *p*< 0.03).

**Figure 3.**
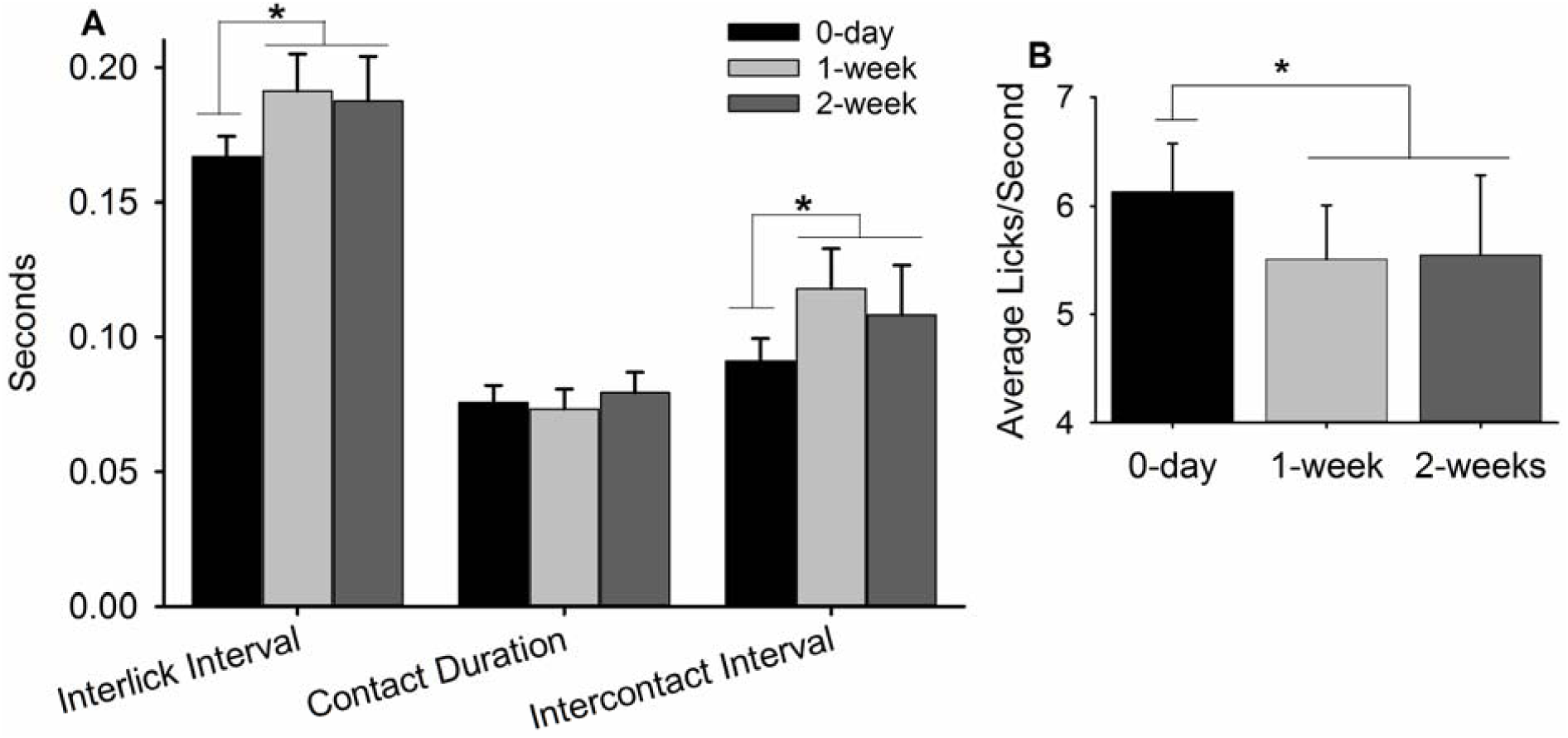
Alterations in the licking microstructure revealed a delay in tongue movement after injury to the mylohyoid muscle. (A) Significant increases in interlick intervals and intercontact intervals were found 1- and 2-weeks post-injury compared to pre-injury. (B) Significant decreases in lick frequency were also noted 1- and 2-weeks after injury. Statistical significance of *p* < 0.05 is indicated by *.

**Figure 4.**
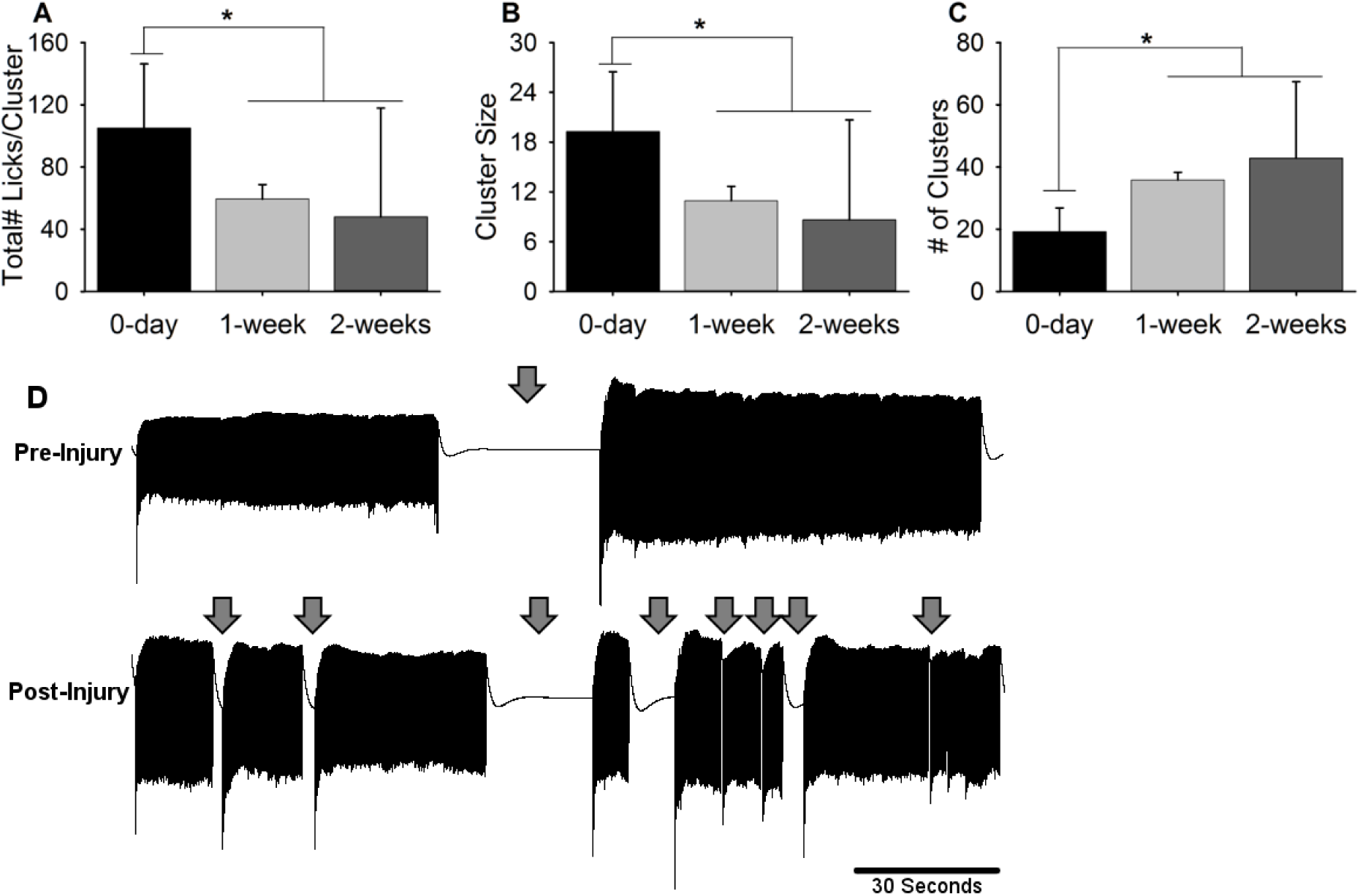
Injury to the mylohyoid influenced the temporal distribution of licking clusters. The rats’ licking behavior was recorded during a 10-minute period of self-drinking. A cluster was defined as a group of sequential licks separated by intervals >500ms. (A) Significant decreases in the total number of licks per cluster were found 1- and 2-weeks post-injury compared to pre-injury. (B) Cluster size, which is the average duration of each cluster, also decreased post-injury. (C) Lastly, there were significant increases in the number of clusters performed post-injury compared to pre-injury. Statistical significance of *p* < 0.05 is indicated by *. (D) An example plot showing the distribution of licking clusters within a 3-minute window pre-injury and post-injury. Arrows denote pauses between clusters of licking. Note the presence of shorter clusters post-injury that occur more frequently. These results suggest that mylohyoid injury modifies both cluster number and duration, provoking repeated pauses/breaks between licking.

Mylohyoid injury also had a significant effect on licking cluster. The duration of each cluster (*F*(2,10) = 4.47, *p* = 0.04) and the number of clusters initiated (*F*(2,10) = 4.27, *p* = 0.04) during the drinking session were affected by injury. Shorter periods of continuous licking were found after injury compared to pre-injury (1 week: *p*< 0.05, 2 weeks: *p*< 0.02) based on the reductions found in cluster size. A significantly higher number of clusters were initiated 2-weeks post-injury compared to pre-injury (*p*= 0.002). The total number of licks per cluster was also evaluated, which is the number of continuous licks between two breaks in licking activity. Significant decreases were found (*F*(2,10) = 4.69, *p* = 0.04) in licking rate per cluster after injury compared to pre-injury (1-week: *p*< 0.03; 2-weeks: *p*< 0.02).

### Depression in Swallowing Motor Activity after Mylohyoid Injury

The mylohyoid (*F*(1,2) = 83.08, *p* = 0.01) and thyropharyngeus (*F*(1,2) = 473.67, *p* = 0.01) muscles were found to have significant differences with EMG amplitudes (% of maximum; Figure 5). Injury resulted in decreases in mylohyoid (*p*< 0.02) and thyropharyngeus (*p*< 0.002) amplitude during swallowing compared to pre-injury. No significant effects of injury were found with the geniohyoid (*F*(1,2) = 8.46, *p* = 0.10), thyrohyoid (*F*(1,2) = 16.24, *p* = 0.06), and thyroarytenoid (*F*(1,2) = 3.63, *p* = 0.20) muscles.

**Figure 5.**
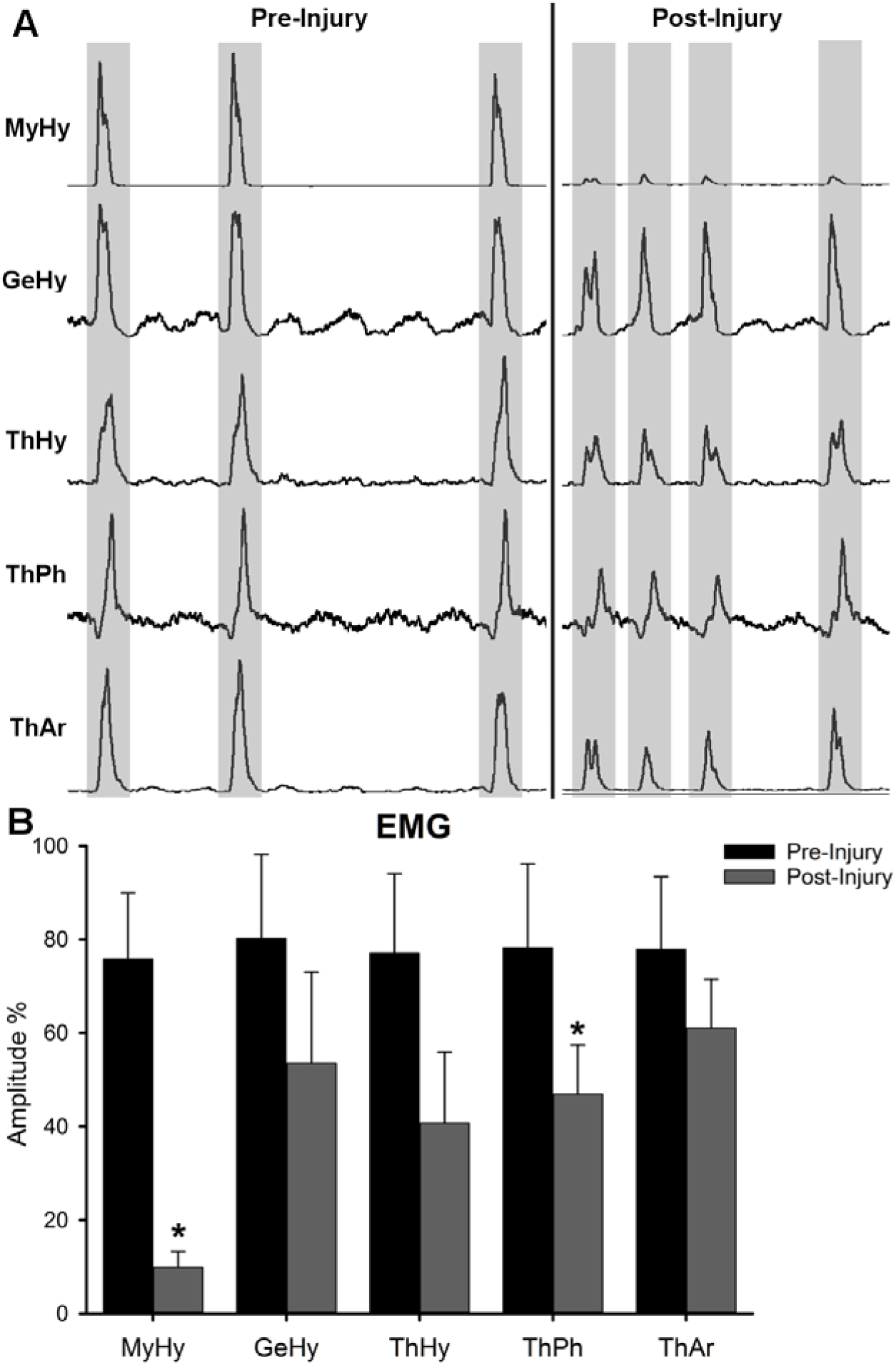
(A) Rectified and integrated EMG activity from the mylohyoid (MyHy), geniohyoid (GeHy), thyrohyoid (ThHy), thyropharyngeus (ThPh), and thyroarytenoid (ThAr) muscles during swallowing pre- and post-injury. Swallowing was identified by MyHy and ThHy activity. Note the decrease in the burst amplitude of each muscle post-injury. (B) Results demonstrating the mean and standard deviations for EMG amplitude (% maximum) of each of the muscles tested. Significant decreases in MyHy and ThPh activity during swallowing were found post-injury compared to pre-injury. Statistical significance of *p* < 0.05 is indicated by *.

## Discussion

We investigated the effects of oral and pharyngeal swallowing function after cryoinjury to the mylohyoid. Similar to other models [28,31], cryoinjury is a feasible and reliable method for creating fibrosis. The cryoinjury increased accumulation of collagen bilaterally in the mylohyoid muscle by 2-weeks post-injury. However, structural damage only partly explains swallowing dysfunction following muscle injury, as abrupt alterations in muscle activity were also evident. Together, these injuries were found to reduce tongue movement and alter bolus parameters during swallowing. Our results support the hypothesis that injury to a single, important swallow-related muscle produces adaptations in oropharyngeal swallowing to compensate for sensorimotor deficits.

Changes in muscle structure, i.e. fibrosis, play a major role in its mechanical properties; however, our results do not confirm our hypothesis that increases in collagen content in the mylohyoid corresponds with alterations in swallowing behaviors. This is because both videofluoroscopy and lick testing demonstrated persistent declines in the licking rate during drinking, starting prior to development of granulation tissue at one-week post-injury. Furthermore, immediate decreases in mylohyoid and thyropharyngeus EMG motor activity during swallowing were observed after injury. Previous work has shown extensive infiltration of inflammatory cells 5-7-days after cryolesion to the mouse heart [28] or rat hindlimb muscle [44], which subsides within 10-14 days, leaving scar tissue to remain. As such, it is possible that swallowing function was negatively impacted by the initial inflammatory response provoked by cryoinjury to the mylohyoid, thus providing evidence for time-specific effects on swallowing.

Since the tongue and jaw move simultaneously during repetitive licking in the rat [45], we can indirectly measure lick frequency via fluoroscopy by calculating the jaw excursion rate [36] and compare this to bolus transit. Along with reductions in licking frequency, we detected alterations in bolus volume within the pharynx. Bolus size is an indirect estimate of bolus volume [46]. In particular, we found an inverse relationship between bolus size and bolus speed through the pharynx one week after injury. Although bolus speed returned to normal levels two weeks after injury, bolus size during swallowing remained small (−35% from baseline). One possible explanation for this is that the rats are compensating for a sensorimotor deficit by reducing their bolus volume. When liquid volume during swallowing decreases, less pressure is needed for bolus transport through the pharynx and the relative timing of swallowing events shorten [47-49]; thus reducing the effort required for swallowing. Previous work in an infant pig model of recurrent laryngeal nerve lesion, which affects vocal fold closure, compared differences between the bolus area in the vallecula and the onset of penetration or aspiration during swallowing [50]. They found changes in bolus size post-lesion, with smaller boluses correlating with increased airway protection. It is possible that sensory adaptations from the nerve injury impacted the central pattern generator (CPG) located in brainstem leading to difficulties in handling the larger boluses safely. The CPG modulates swallowing movement through sensory feedback. Prior work has shown that sensory input from the oral cavity can influence bolus volume and direct the pharyngeal motor response during swallowing [51]. Unlike pigs, rats are not known to aspirate food/liquid. As such, the alterations in bolus parameters found after mylohyoid injury may reflect a means of overcoming challenges found with oral function in order to maintain bolus transit. Our results are consistent with clinical findings, which have shown that patients self-attenuate the effects of a swallowing deficit by reducing their bolus volume per swallow [52]. In further support of a compensatory strategy taking place, a decrease in the frequency of swallows occurred concurrently with the increase in bolus speed one-week post-injury. The observed changes may relate to alterations in oropharyngeal movement. Swallowing frequency was determined by dividing the total number of swallows by the time between the start and termination of >3 sequential swallows. Thus, the time accounts for the individual swallowing act and the interval between consecutive swallows. However, the latter was not shown to be affected by injury. To characterize the full extent of differences in functional ability that result from mylohyoid injury, further analysis of swallowing kinematics is warranted.

Further characterization of the licking pattern demonstrates that the injury provoked increases in interlick intervals, which indicates that tongue protrusion and retraction were prolonged during drinking. Changes in interlick intervals have also been attributed to delays in swallowing movement [38]. However, we cannot substantiate this with our study because of the limited visualization of the hyolaryngeal structures with our fluoroscopy equipment. Of note, impaired base of tongue retraction during swallowing is a common problem in patients treated with radiation for oropharyngeal cancers [53]. Tongue base retraction is critical for effective bolus clearance through the pharynx during swallowing [54]. These impairments are often attributed to high doses of radiation to tongue or pharyngeal muscles [55,54]; however, our results demonstrate that injuries to the mylohyoid muscle can also reduce tongue displacement during swallowing.

The total number of licks per drinking session did not differ with injury, suggesting that similar total volumes of liquid were consumed during each session. However, the way in which liquid was consumed after injury was very different. A licking cluster is a sustained period of licking terminated by breaks in licking, i.e., when the head backs away from the spout; its summation reflects the time spent drinking. Licking clusters in normal rats consisted of long periods of continuous licking (∼19.27 seconds) for about ¾ of the drinking session. After injury, we observed short-bursts of uninterrupted licking for ∼8-11 seconds in duration that occurred repetitively throughout the session. As a result, the number of clusters initiated increased by ∼50% post-injury. These observations suggest that injured rats took repeated breaks during the drinking session and required longer meal times to consume the same volume of liquid compared to pre-injury. Clinically, lengthy feeding times can lead to fatigue and put patients at risk for developing malnutrition [56]. Rats are known to fluctuate the duration of time spent licking when the temporal pattern of licking is altered in order to maintain stable liquid consumption [40]. Previous studies in normal rats have indicated that with palatable solutions the cluster duration is dependent on licking rate and interlick interval duration. For example, the lick rate and cluster duration is enhanced in rodents with consumption of a familiar flavor of a pleasant solution (e.g., sucrose) [57]. With non-palatable solutions (e.g., ethanol), decreases in licking rate have been shown to result in longer sustained bursts [39]. In our experiments, rats were given chocolate milk, which is a palatable solution and may have unintentionally positively reinforced them to drink for longer periods. However, contrary to the above findings, after injury we detected an increase in interlick interval duration during shorter periods of sustained licking. Thus, injured rats took longer to move their tongue and they required frequent breaks during licking, regardless of their motivation. A possible cause that warrants further exploration is that mylohyoid injury is inhibiting the strength and endurance required to perform prolonged, sustained periods of licking.

There is evidence implicating that the effects of muscle injury on movement dysfunction are partly attributed to sensory changes [58,59]. In an initial attempt to address this question, we measured the motor activity of five important swallowing-related muscles after injury to the mylohyoid. The swallowing CPG initiates and coordinates motor action via sensory input from peripheral nerves. As such, EMG testing provides important information on the motor response of synergistic muscles during swallowing and the potential central mechanisms. If changes in the EMG amplitude were only observed in the mylohyoid muscle, which was injured, then swallowing dysfunction is likely attributed to a decline in motor function. However, our data revealed that not only was mylohyoid activity reduced during swallowing, but responsiveness of the thyropharyngeus muscle was also reduced, which was not directly injured. Anatomical evidence has reported an extensive convergence of trigeminal and vagal afferent nerves in the brainstem [60], which would facilitate the expansion of the sensory receptive fields after injury [61]. Inflammation in the facial and lingual muscles have been shown to inhibit swallowing reflexes [26,27]. Further, when injury or inflammation is experimentally induced in a hindlimb muscle, studies have demonstrated that activity of the injured muscle is reduced and redirected to synergistic muscles [58,59]. Therefore, injury to the mylohyoid may compromise swallowing via sensory disturbance.

Our study has several limitations. Our sample size was insufficient for detecting small/medium effects in EMG data. Although a depression in EMG amplitudes was observed across all the muscles tested, larger sample sizes are needed to study these differences. Sample size calculations were based on changes in the mylohyoid muscle. Similar studies in other animal models have indicated that three animals are sufficient at detecting differences with EMG using a within-subject design, which increases statistical power and reduces the sample size needed [62]. Furthermore, no sham-surgical control animals were included to determine if surgery itself impacted licking and swallowing function. The incision design (i.e. location at midline and short size) was chosen to limit the extent of skin injury, resulting in complete closer of the wound within 3-4 days post-surgery.

## Conclusion

This is the first investigation analyzing the effects of a single muscle injury on swallowing function. Disruptions in oral and pharyngeal swallowing were detected, including delays in tongue movement and alterations in bolus flow. Findings demonstrate that changes in swallowing dysfunction are occurring prior to the onset of tissue level changes (i.e., fibrosis or inflammation). These results offer initial information about swallowing dysfunction in response to muscle injury. Further analysis with other muscles of interest to swallowing are needed to improve our understanding of the effects of injury-induced adaptations.

### Ethical approval

All applicable international, national, and/or institutional guidelines for the care and use of animals were followed.

